# Shared Neuroanatomy, Separate Mechanism: *in vivo* ERK and mTOR Manipulations Reveal Female-Specific Molecular Signaling for Auditory Forebrain-Dependent Learning in Juveniles

**DOI:** 10.64898/2026.07.02.736152

**Authors:** Kruthika V Maheshwar, Siddhanth Chari, Sarah E London

**Affiliations:** Department of Psychology, The University of Chicago; Institute for Mind and Biology, The University of Chicago; The College at The University of Chicago; Neuroscience Institute, The University of Chicago

## Abstract

Developmental experience can produce lasting changes in neural function and behavior. Zebra finch offers a powerful model for identifying the molecular mechanisms underlying this process. Both juvenile male and female zebra finches perform developmental sensory song learning that influences their adult behaviors: in males, the structure of the song they sing and in female, the song preferences they exhibit (females cannot sing). The auditory forebrain, a region distinct from but connected to nodes of the male singing circuitry, is required for male sensory song learning. Song experience induces epigenetic, genomic, molecular, cellular and systems-level alterations in the auditory forebrain of males. Much less evidence is available for females. Although epigenetic and molecular data implicate the auditory forebrain in female sensory song learning, there has been no causal test of its role. Further, molecular evidence indicates the potential for distinct mechanisms for male and female sensory song learning, even though they learn during a largely overlapping developmental period. We used pharmacological manipulations of the ERK and mTOR cascades in the auditory forebrain of juvenile females during controlled tutoring, and an operant assay for adult song preference, to test the causal role of the auditory forebrain and the two cascades known to be required for male sensory song learning. We demonstrate that the auditory forebrain is required for female sensory song learning, and that while ERK signaling is necessary for both sexes, that of mTOR is sex specific. Results raise implications for alternative molecular cascade cross-talks and protein synthesis processes that successfully support the developmental learning at the same age and brain region.

## INTRODUCTION

The developing brain converts early experiences into lasting changes in neural function and behavior^1–4^. Zebra finches provide a precise and tractable model for investigating the molecular mechanisms underlying this process. Exposure to song during development has lifelong consequences for behavior in these birds^4–6^. In males, exposure to a “tutor” song during the Critical Period (CP) for sensory song learning, from posthatch day (P) 30-65, guides song production in adulthood^4,7–9^. While females do not sing, they also undergo developmental sensory song learning, too. Adult females exhibit preference to songs they experienced during development^6,10,11^. Identifying the molecular mechanisms by which early song experience is learned can provide insights into how developmental experiences shape the brain and behavior more generally.

Much of what is known about how the brain encodes early song experiences comes from work in males. In males, the auditory forebrain lobule (AL) is necessary for developmental song learning^12^. Regulated activation of two pathways — the extracellular signal-regulated kinase (ERK) and mechanistic target of rapamycin (mTOR) — is necessary during tutor experience for high fidelity tutor song copying^12,13^. Whether or not females require the same neural and molecular mechanisms for developmental sensory song learning is less clear. The crucial test for a canonical CP – the extension of learning ability with isolation from relevant experience – has not been performed in females. Behavioral tests for song preference learning, however, indicate that female sensory song learning occurs ∼P20-70, overlapping but extended with males^7,14,15^. Immediate early genes, including those regulated by ERK, are rapidly transcribed in juvenile female AL after song experience^16,17^. Yet song does not increase phosphorylated ribosomal protein S6 (pS6), a marker of mTOR activity, in female AL at P23 or P30, ages at which behavioral data indicate successful song preference learning^13,18^. Notably, pS6 is also a marker for activity-dependent translation; new protein synthesis is a conserved mechanism of long-term memory formation^19^. Females may thus also rely on the AL for sensory song learning, but through distinct molecular pathways than males.

Here, we tested this postulation by combining controlled tutor experience with *in vivo* manipulations of ERK and mTOR signaling cascades targeted specifically to the AL in juvenile females. We used the same experimental strategy as in males to facilitate direct comparisons^12,13^, but instead assessed adult song preference using an operant perch hop song assay. Our results establish that juvenile sensory song learning requires the AL in females, as it does in males, but via distinct mechanisms.

## MATERIALS AND METHODS

### Experimental Animals

All procedures were conducted in accordance with the NIH guidelines for the care and use of animals for experimentation and were approved by the University of Chicago Institutional Animal Care and Use Committee (ACUP no.72220). All birds were housed on a 14:10 hour light: dark cycle with seed and water being provided *ad libitum*. Juveniles used for the experiment hatched in flight aviaries and lived there among males and females of all ages. At P23, they were removed from the aviary to prevent song exposure other than from tutor sessions (**Fig. 1**). They were then moved into acoustic chambers with two foster females and up to 3 age matched siblings/ foster siblings to provide a near-normal social environment. On day 30, each juvenile female was moved to be housed with a single female, and this living condition was maintained for the duration of the study.

**Figure 1.**
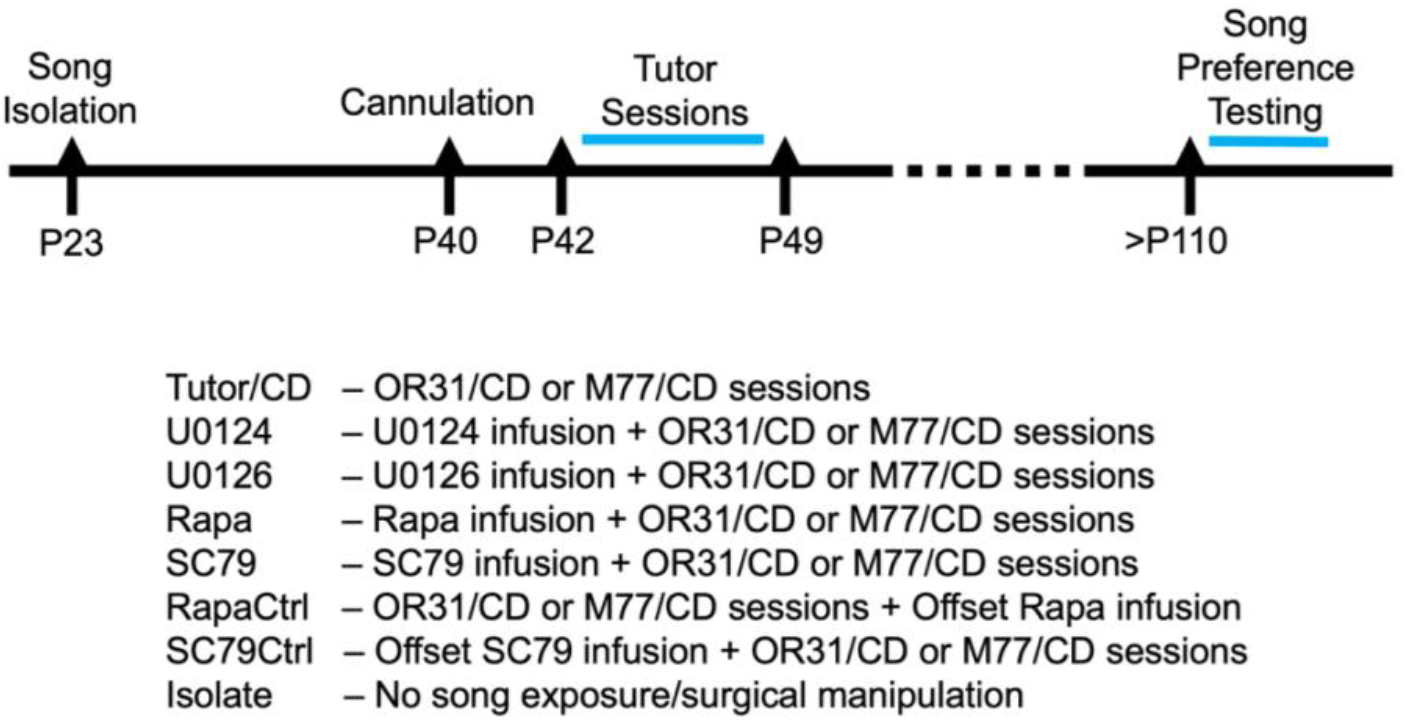
Schematic Representation of Experimental Timeline and Groups. Song isolation for the experimental birds started at P23. All tutor exposure occurred from P42-P49. Cannulation surgeries for all groups except the Tutor/CD and Isolates was carried out on P40. U0124, U0126, Rapa, and SC79 groups received infusions 30 minutes prior to tutor sessions; the RapaCtrl and SC79 birds received infusions offset from tutor sessions. Preference testing was carried out at adulthood for six consecutive days.

### Surgery

All groups except the Tutor/CD and Isolate groups underwent surgery. At P40 (**Fig.1**), the birds in these groups were implanted with a bilateral cannula with an intercannulae distance of 1mm and length on 2mm into their AL; coordinates 70 mm anterior to Y0 (the anterior-most boundary of the cerebellum at midline) 50 mm lateral to midline, at a head angle of 35 °, under monitored isoflurane anesthesia^12^. Birds awoke 2-5 mins post-surgery after which a sharp noise was made on either side of the ear to ensure that hearing was still intact.

### Tutor Sessions

We followed a previously established paradigm^12,13^. Two tutors — OR81 and M77—were used with birds in each group being assigned a tutor randomly such that 50% of the birds in each group experience OR81 and his song and 50% experienced M77 and his song. All experimental birds experienced eight tutor sessions, one per day from P42 to P49 (**Fig. 1**). They received four sessions in the first 7 hour of lights-on (except the first hour after the lights are on; “morning session”), and four in the second 7 hour of lights on (except the last hour of the day; “afternoon session”).The tutor session consisted of a combination of pre-recorded songs as well as live tutor exposure for 1.5 hours^12,13^.

### Experimental groups and Drug Infusions

Birds were assigned to one of eight groups (n = 8/group; **Fig. 1**). Tutor/CD birds received tutor song exposure with no surgical or pharmacological manipulation; Isolate birds received no song exposure, staying with their female companions throughout. The remaining six groups received tutor exposure paired with infusions targeting ERK or mTOR signaling in the AL. U0126 was the ERK inhibition group and the U0124 group acted as the control. Rapamycin (Rapa) and SC79 groups were the mTOR manipulation groups — Rapamycin inhibits mTOR directly, while SC79 constitutively activates AKT, upstream of mTOR. Birds in the U0126, U0124, Rapa, and SC79, groups were infused with 0.5 µl of U0126 (20ug/µl), and U0124 (20µg/µl), rapamycin (1µg/µl), and SC79 (200ng/µl) respectively, 30 mins prior to the start of each tutor session, as we had previously validated^12,13^. The Rapamycin offset control group (RapaCtrl) received rapamycin infusions four hours after the end of the tutor session^13^. The SC79 offset control group (SC79Ctrl) received SC79 infusions either 2 hours after the end of the morning session or 2 hours before the start of the afternoon session^13^.

### Hot Zone Scoring

To acquire a metric of social engagement during tutor sessions, we defined a subsection of the cage housing the juvenile and the foster mother as the “Hot Zone” (HZ)^13^. The HZ was the third of the juvenile’s cage closest to the tutor bird cage. It extended throughout the dimensions of the cage, from floor to ceiling and wall to wall, and was marked by the position of a cage door/perch. A scorer blind to experimental groups used JWatcher^20^ to quantify the time the juvenile spent inside the HZ attending to the tutor (looks). Three tutor sessions (day 1,4, and 8) were scored per bird.

### Perch Trigger Assay

Song preference in adult females was assessed using an operant perch-trigger assay. The test apparatus was placed on the middle tier of a three-level wire rack and consisted of a 17 × 12 × 14-inch cage equipped with four landing perches. Balsa wood dowel perches (⅛ inch) were positioned 8 inches above the cage floor, with the pair on the same wall separated by 5 inches, and the opposing pair slightly staggered. Song stimuli were presented through speakers placed just outside the cage, directly behind the song-associated perches. The birds generally preferred to remain toward the back of the cage, so food and water dishes were positioned at the front to encourage use of the whole space. These dishes were swapped every second day. To provide a social context, the experimental cage was flanked by cages containing a pair of adult females each. These pairs were rotated daily to avoid potential social biases. In addition, two cages housing groups of three females each were placed on the shelf below the experimental cage.

The experimental birds were tested once they reached adulthood. A pair of diagonally positioned perches was designated as the song perches; landings on these perches triggered song playback. The remaining two perches served as control perches, where landings did not elicit playback. All perches were connected to Raspberry Pi devices, which controlled the playback and simultaneously logged every perch hop. Testing was conducted over six consecutive days in two-hour sessions, scheduled either 2–4 hours or 5–7 hours after lights-on. Each bird completed three sessions in each time slot. To control for side biases, the perch–song assignment was reversed on the fourth day. Birds were transferred to the experimental cage 30–45 minutes prior to each session for acclimation and returned to their home chambers at the end of the session.

### Data Analyses

#### Perch preference assay

##### Variance Analyses

To evaluate differences in variability across the eight experimental groups, we analyzed the variance of mean hopping activity. We first performed Levene’s Test for Homogeneity of Variance (based on absolute deviations from the mean) on the mean number of hops per day for each bird to determine if the dispersion of bird-level means differed significantly across the groups. We then followed this up with an analysis to check if the variances between the song hops and the control hops within each of the groups were different. For this, a Levene’s test for equality of variances was performed on the mean number of hops per day for each bird for each group.

##### Perch Type Preference Analyses

To assess perch type preference, we calculated two complementary Perch Preference Ratios: the Song Perch Preference Ratio and the Control Perch Preference Ratio. These were defined as the number of hops on the song or control perches, respectively, normalized by the total number of hops. Consequently, for each bird, six Song Perch Preference Ratios and Control Perch Preference Ratios were obtained, corresponding to the six testing days. Each experimental group consisted of eight birds, thus resulting in eight sets of six ratio pairs per group. Within-group differences in perch activity were evaluated using a single-factor Linear Mixed-Effects Model (LMM). The two Preference Ratios served as the outcome variable, modeled as a function of PerchType (Song vs. Control) as a fixed effect. To account for the dependency of repeated measures across the six testing days, Bird ID was included as a random intercept. The model was expressed as:

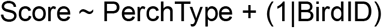

Population-level effects were visualized by extracting Estimated Marginal Means (EMMs) and their corresponding 95% Confidence Intervals (CIs).

##### Song Type Preference Analyses

1. Groupwise preference test To check if the tutored groups and the Isolate group formed a song preference, we calculated two ratios: the Tutor Song Preference Ratio and the Unfamiliar Song Preference Ratio. Each ratio was defined by dividing the number of hops on the respective song perch (Tutor or Unfamiliar) by the total number of hops. Thus, for each bird, six Tutor Song Preference Ratios and Unfamiliar Song Preference Ratios were obtained corresponding to the six testing days. Each experimental group consisted of eight birds, thus resulting in eight sets of six ratio pairs per group. Within-group differences in song type preferences were evaluated using a single-factor Linear Mixed-Effects Model (LME). The two Preference Ratios served as the outcome variable, modeled as a function of SongType (Tutor vs. Unfamiliar) as a fixed effect. Bird ID was included as a random intercept. The model was expressed as:

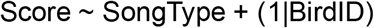

Population-level effects were visualized by extracting Estimated Marginal Means (EMMs) and their corresponding 95% Confidence Intervals (CIs).
2. Between group comparisons To compare the strength of song preference across our experimental groups, we first defined a metric called the Preference Index. Preference Index was obtained by dividing the difference between tutor hops and unfamiliar hops by the total number of song hops. This index ranged from -1 to 1, with indices less than 0 indicating unfamiliar song preference and indices above 0 indicating preference to the tutor song. For each bird, this generated 6 indices corresponding to the 6 test days, and each group thus has 8 sets of six indices. To account for this nested structure, we employed a hierarchical (multi-level) bootstrapping procedure with 1,000 iterations. We used a two-stage resampling strategy. For each iteration, eight birds were sampled with replacement from the cohort.

For each selected bird, six daily observations were sampled with replacement from that individual’s data pool. The mean for each bird was calculated, and these were then averaged to yield a single group-level bootstrap estimate per iteration. This process resulted in a distribution of 1,000 bootstrapped means for each of the eight experimental groups. To do the pairwise bootstrapped contrasts, for every iteration, the mean of the second group was subtracted from the mean of the first group to generate a distribution of the differences (Δ). We quantified the likelihood of the observed effect by calculating the Probability Mass > 0, representing the percentage of bootstrap iterations where the mean of the first group exceeded that of the second. To visualize the pairwise comparisons we generated Probability Density Functions (PDFs) for each pairwise contrast. Each histogram represents the frequency of the 1,000 bootstrapped mean differences, with a zero-reference line denoting the null hypothesis and a central tendency line indicating the mean of the difference distribution.

#### Hot Zone Analyses

As a measure of the juvenile’s social engagement with the tutor, we investigated potential differences between the groups in the time spent inside the HZ engaging with the tutor (looks). We defined LookTimes as the proportion of the TS that the juvenile spent looking at the tutor. LookTimes were analyzed using a Linear Mixed-Effects Model (LME) to accommodate the unbalanced experimental design (unequal number of subjects across the eight cohorts). We defined Group (8 levels) and Day (3 levels) as interacting fixed effects. To account for the dependency of repeated observations within the same individual, a random intercept for BirdID was included.

## RESULTS

### While the hopping activity was variable across groups, all groups except the Isolate group showed comparable variances between the song and control perches

To check if variation in hopping was evenly distributed across the groups, we first performed a Levene’s test for equality of variances on the absolute number of hops per day for each bird (**Fig. 2**). The Levene’s test was significant (F_7, 55_ = 2.64, p = 0.020), indicating that the variances in mean hopping were not equal across groups.

**Figure 2:**
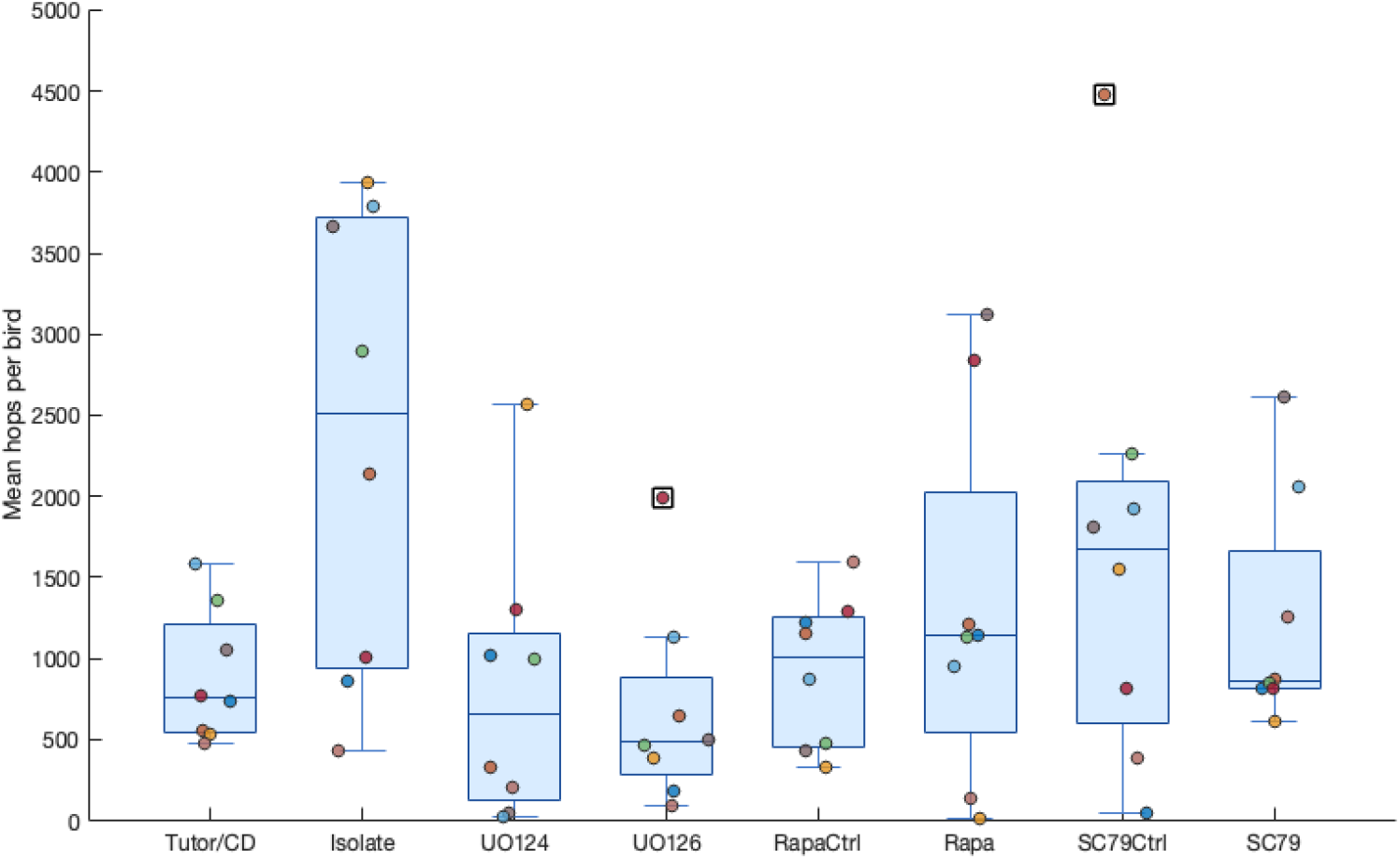
Birds across groups showed unequal variances. Box and whisker plot showing the spread of mean number of hops per bird across the eight groups. Box edges represent the interquartile range (IQR), the horizontal line inside each box marks the median, and whiskers extend to the most extreme data points within 1.5×IQR of the box edges. Colored circles represent the mean hop value for each individual bird within a group. Black squares (SC79Ctrl y∼4500 and U0126 y∼2000) indicate outliers, defined as values falling beyond 1.5×IQR from the box edges. Individual dots represent individual birds.

Follow-up F-tests comparing group variances revealed that the Isolate group had significantly higher variance than the Tutor/CD, RapaCtrl, and the U0126 (p= 0.0038, 0.0085, 0.041). The SC79Ctrl group also had significantly greater variance than the Tutor/CD and the RapaCtrl groups (p = 0.0046,0.010). Additionally significant differences were observed between the Rapa and Tutor/CD groups (p = 0.016), and the Rapa and RapaCtrl groups (p = 0.03). We then performed a groupwise Levene’s test for equality of variances on the absolute number of hops on the song perches and control perches. We found that the variances between the song perches and control perches were not significantly different for any of the groups except the Isolate group, which showed a significantly higher variance on the song perches relative to the control perches (p = 0.00019706).

### Birds across groups prefer song perches significantly over the control perches

Prior work showed that song serves as a reward^6^. To validate our assay, we examined the PerchType preference of birds across the groups by using a Linear Mixed-Effects Model to compare the Song Perch Preference Ratios (defined as the number of hops on the song perches normalized to total number of hops; i.e.; song hops + control hops) and the Control Perch Preference Ratios (defined as the number of hops on the control perches normalized to total number of hops).

We found that all groups preferred the song perches significantly over the control perches (**Fig. 3**): Tutor/CD (LMM: F_1,94_=15.03, p=1.96×10^-4^. Mean ± SEM: song 0.555 ± 0.020, control 0.445 ± 0.020); Isolates (F_1,94_=471.92, p=2.04×10^-38^ : song 0.785 ± 0.019, control 0.215 ± 0.019) ; U0124 (F_1,92_=50.11, p=2.83×10^-10^: song 0.666 ± 0.034; control 0.349 ± 0.033); U0126 (F_1,94_=111.17, p=1.29×10^-17^: song 0.761 ± 0.030 control 0.285 ± 0.034; RapaCtrl (F_1,94_=29.40, p=4.55×10^-7^: song 0.597 ± 0.026, control 0.403 ± ± 0.026); Rapa (F_1,94_=30.04, p=3.54×10^-7^: song 0.635 ± 0.035, control 0.381 ± 0.035); SC79Ctrl (F_1,94_=33.58, p=9.07×10^-8^: song 0.618 ± 0.029, control 0.382 ± 0.029); SC79 (F_1,94_=15.96, p=1.29×10^-4^: song 0.582 ± 0.029, control 0.418 ± 0.029 respectively).Importantly, significant preference for the song perches was observed even in the Isolate group, indicating that song served as a rewarding stimulus independent of developmental song exposure.

**Figure 3.**
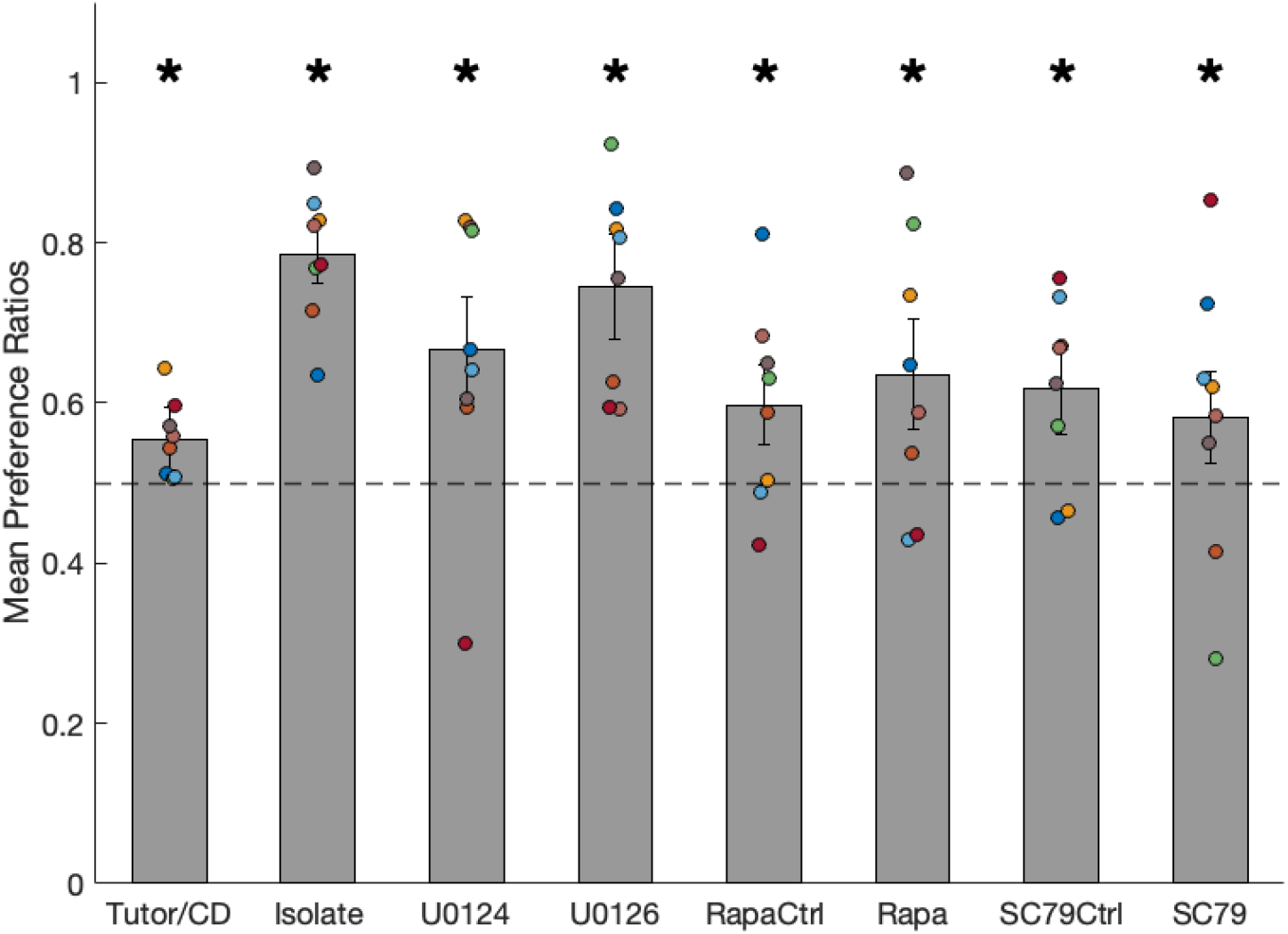
Birds across all groups preferred the song perches significantly over the control perches. Means of Song Perch Preference Ratios (Song Perch Preference Ratio = Song Hops/Total Number of Hops) for each group. Error bars indicate SEMs and (*) indicate significance above chance (dashed line). Colored dots represent individual birds.

### The Tutor/CD group showed a preference for the tutor song, demonstrating developmental sensory song learning. Isolate group showed no consistent song preference

We next assessed the presence of SongType preference in each of the groups by using LMM to compare the Tutor Song Preference Ratios (defined as the number of hops on the tutor song perches normalized to total number of hops; i.e.; song hops+ control hops) and Unfamiliar Song Preference Ratios (defined as the number of hops on the unfamiliar song perches normalized to total number of hops).

Unmanipulated adult female zebra finches prefer songs they were developmentally exposed to over an unfamiliar match^6^. In line with this, the Tutor/CD group preferred the tutor song significantly over the unfamiliar song (LMM: F_1,94_=17.52, p=6.41×10^-5^. Mean ± SEM: tutor 0.341 ± 0.024, unfamiliar 0.214 ± 0.019) (**Fig. 4**), demonstrating evidence for developmental sensory song learning. We expected that the Isolate group would not show a significant preference to either of the songs^6^, reflecting the absence of developmental song exposure. Consistent with this, the Isolate group did not show a SongType preference ( F_1,94_=0.006, p=0.937: tutor 0.403 ± 0.032, unfamiliar 0.408 ± 0.036) (**Fig. 4**).

**Figure 4.**
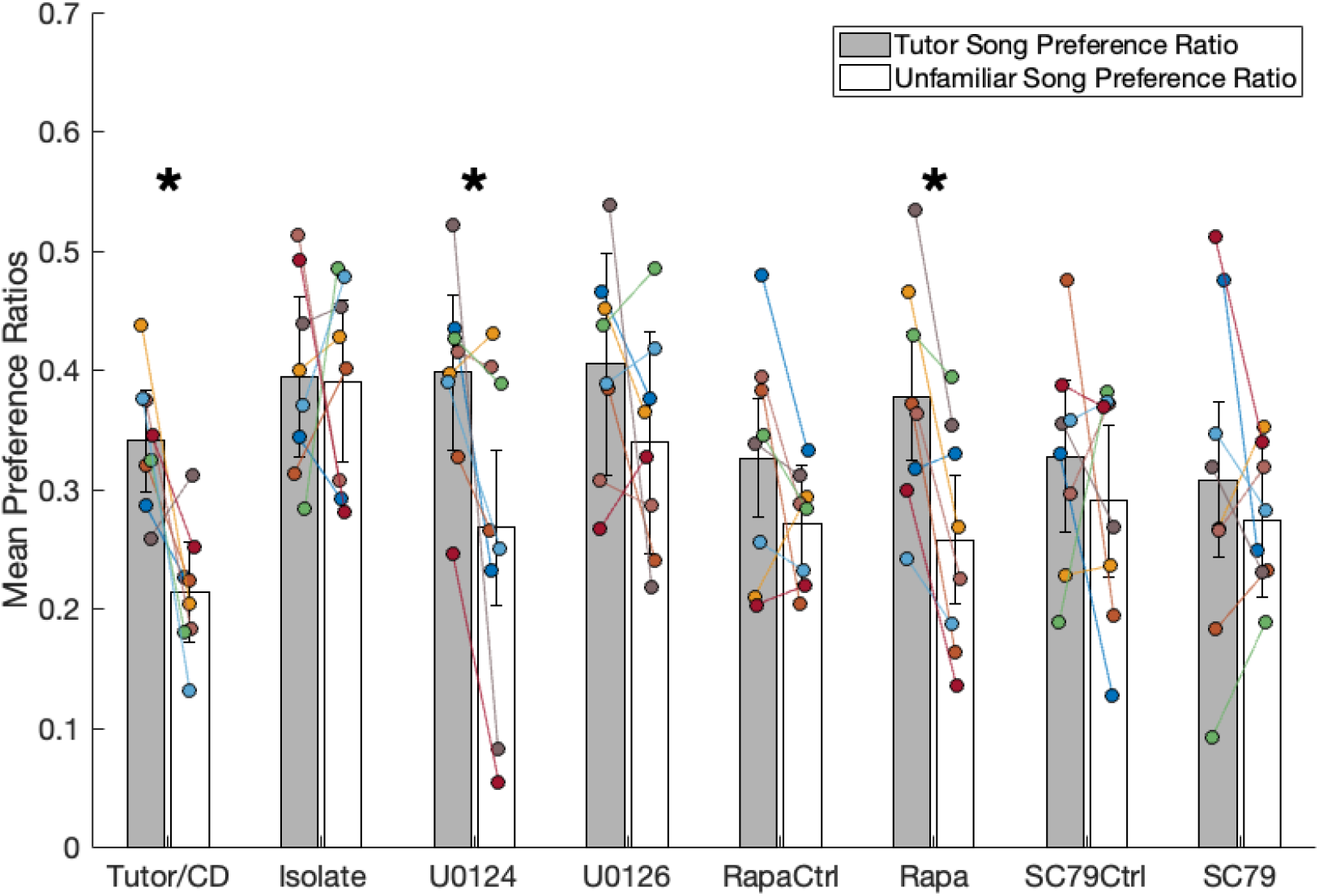
The Tutor/CD group, Rapamycin group, and the U0124 groups significantly preferred the tutor song over the unfamiliar song. The other groups did not show a significant SongType preference. Means of Tutor Song Preference Ratios (Tutor Song Preference Ratio = Tutor Song Hops/Total Number of Hops) and Unfamiliar Song Preference Ratios (Unfamiliar Song Preference Ratio = Unfamiliar Song Hops/Total Number of Hops) for each group. Error bars indicate SEMs and (*) indicate significant difference between Tutor Song Preference Ratio and Unfamiliar Song Preference Ratio. Colored dots represent individual birds.

### Transient suppression of ERK signaling during tutor song experience, bidirectional manipulation of mTOR offset from tutor song experience, and constitutive activation of the mTOR cascade during tutor song experience disrupted song preference formation

The U0126 group (F_1,94_ = 0.98, p = 0.326: tutor 0.432 ± 0.048 unfamiliar 0.362 ± 0.049), the RapaCtrl group (F_1,94_ = 2.44, p = 0.121: tutor 0.326 ± 0.026, unfamiliar 0.271 ± 0.025), the SC79Ctrl group (F_1,94_ = 0.68, p = 0.413: tutor 0.357 ± 0.032, unfamiliar 0.303 ± 0.033), the SC79 group (F_1,94_ = 1.20, p = 0.276: tutor 0.308 ± 0.026, unfamiliar 0.274 ± 0.021) did not show a significant SongType preference (**Fig. 4**).

### The U0124 and the Rapamycin groups showed evidence of having learnt the tutor song

The U0124 group (F_1,94_ = 7.88, p = 0.0061: tutor 0.398 ± 0.033, unfamiliar 0.274 ± 0.034) and the Rapa group (F_1,94_ = 9.80, p = 0.0023: tutor 0.394 ± 0.029, unfamiliar 0.281 ± 0.025) showed a significant preference to tutor song over the unfamiliar song (**Fig. 4**).

### The preference phenotypes of the U0124 and the Rapamycin groups were comparable to that of the Tutor/CD group, whereas those of the U0126, Rapamycin Control, SC79 Control, and SC79 groups were comparable to the Isolate group

To validate the results of our group-level analyses, account for the nested structure of our data, and to carry out between-group comparisons, we conducted a secondary analysis using a hierarchical bootstrap. We used this approach to compare the preference distributions of our experimental groups against the Tutor/CD and the Isolate groups (**Fig. 5A-5M**). We also compared the U0124-U0126, Rapa-RapaCtrl, and SC79-SC79Ctrl pairs (**Fig. 5N-5P**).

**Figure 5.**
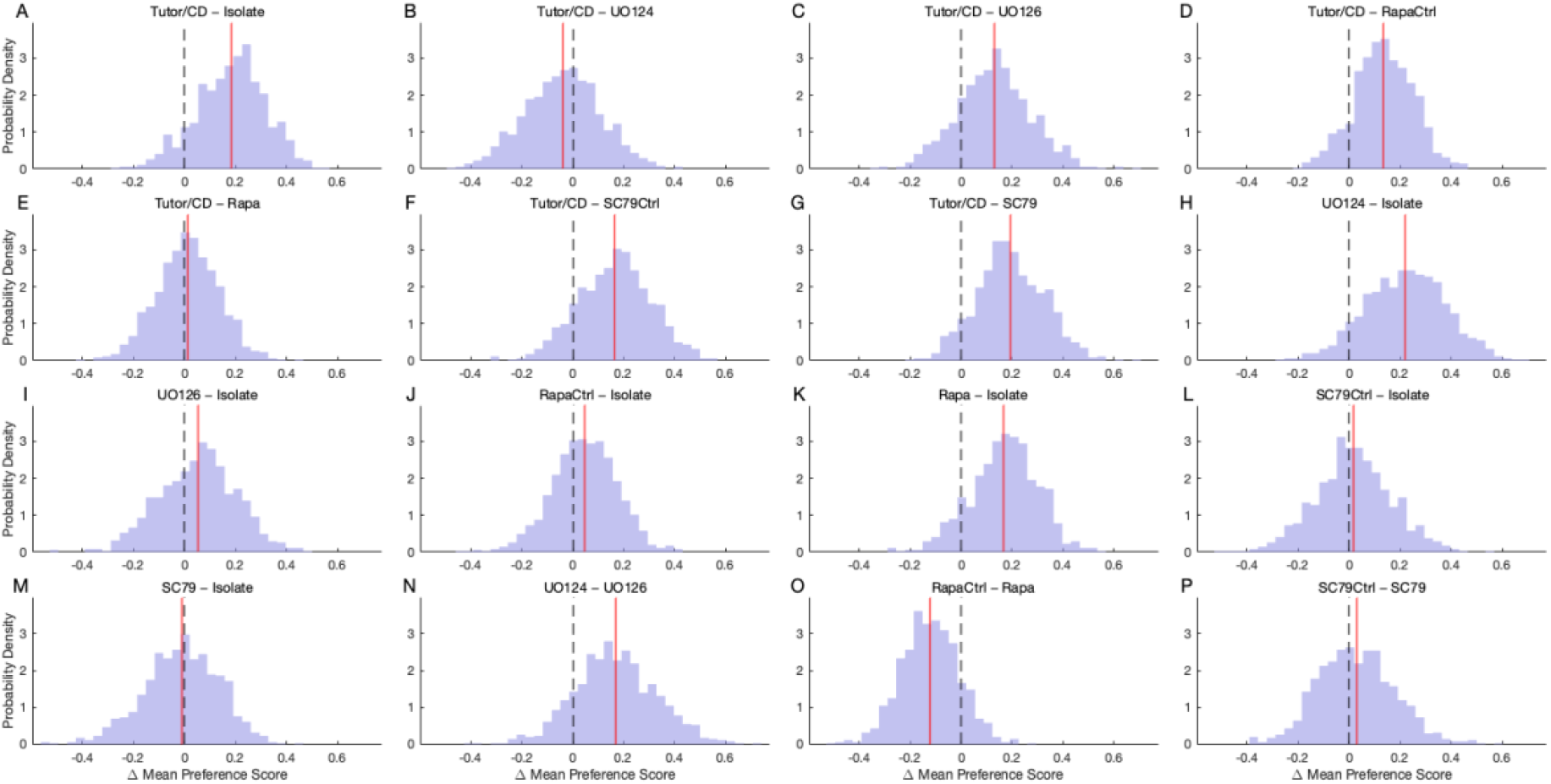
Bootstrap sampling distributions of pairwise mean differences across groups. A-G) Comparisons to the Tutor/CD group. H-M) Comparisons to the Isolate group. N-P) Drug group comparisons. The red line is the mean of the mean difference between the two groups such that if this line is at x>0, the first group has higher preference and if it is at x<0, the second group has higher preference. If the red line and the dotted line overlap, both groups show comparable similarity. The greater the separation between the red and the dashed black line, the more confident the result.

For the Tutor/CD − Isolate comparison, the bootstrap distribution was shifted positively (90.2%>0), supporting a directed preference for the tutor song in the Tutor/CD group (**Fig. 5A**). All other experimental groups segregated into learning and non-learning groups when compared to the Tutor/CD and the Isolate groups. The bootstrapped distribution of the Tutor/CD – U0124 (**Fig. 5B**) comparison means were centered around zero (40.4%>0) suggesting that the birds in the U0124 groups had Preference Scores comparable to that of the Tutor/CD group. Similarly, the distributions of the Tutor/CD – Rapa (55.4%>0) comparison was also centered around zero indicating that the Rapamycin birds were phenotypically comparable to the Tutor/CD birds (**Fig. 5E**).

The U0126, RapaCtrl, SC79Ctrl, and SC79 groups were phenotypically comparable to the Isolate group as evidenced by the U0126 – Isolate (**Fig. 5I**), RapaCtrl – Isolate (**Fig. 5J**), SC79Ctrl – Isolate (**Fig. 5L**), and SC79 – Isolate (**Fig. 5M**) comparisons that were centered around zero (64.1%>0, 65.3%>0, 53.9%>0, and 48%>0 respectively).

### No correlation, global or group-wise, was observed between the activity level and preference indices

It was possible that individual differences in locomotor activity biased our preference indices. We therefore performed a correlation analysis between Total Daily Hops (sum of all activity across song and control perches) and the Mean Preference Index for each bird (n=64 birds).

We found no significant global (r=−0.152, p=0.231; **Fig. 6**) or group-wise (**Table 1**) correlations between the hopping activity and preference indices. This suggests that a bird’s ability to discriminate and prefer a specific song is independent of its activity level and that the preference metrices were not biased by hyperactive or hypoactive birds.

**Figure 6:**
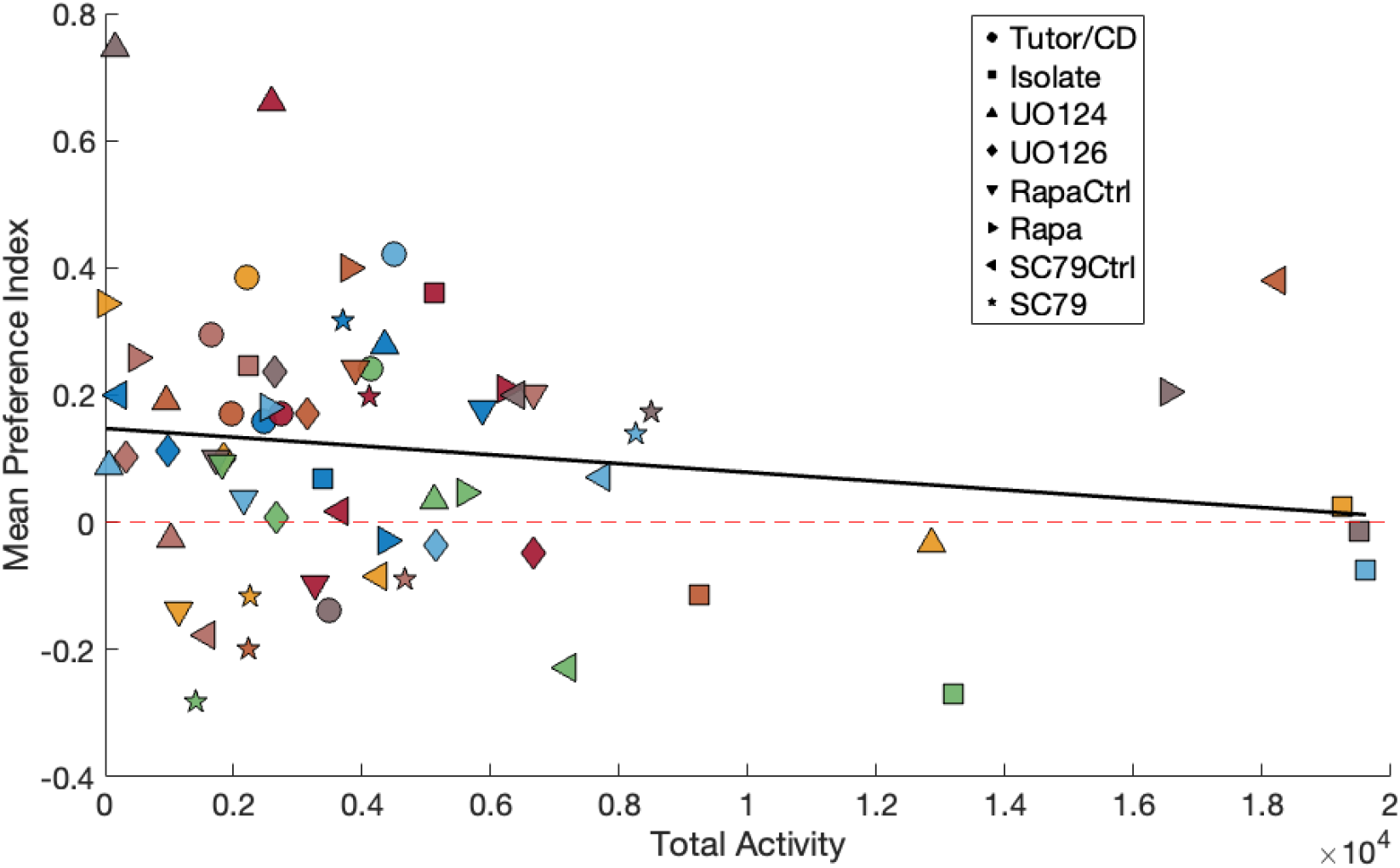
Preference Indices did not correlate with hopping activity. Y axis denoted mean preference index. Individual dots represent individual birds. The black line shows the pooled linear trend across all birds (Pearson’s r = −0.152, p = 0.231), indicating no significant relationship between total activity and mean preference index. The red dashed line marks a preference index of zero.

**Table 1:** Group wise r and p values for correlations between total hops and preference indices.

|  |  |  |
| --- | --- | --- |
| **Group** | **r-value** | **p** |
| Tutor/CD | 0.063 | 0.88 |
| Isolate | -0.63 | 0.09 |
| U0124 | -0.21 | 0.62 |
| U0126 | -0.629 | 0.09 |
| RapaCtrl | 0.45 | 0.26 |
| Rapa | -0.16 | 0.7 |
| SC79Ctrl | 0.436 | 0.28 |
| SC79 | 0.36 | 0.38 |

### HZ analyses showed that the groups did not differ significantly in the time they spent engaging with the tutor

Sensory song learning in male zebra finches is socially mediated^21,22^. However, conflicting evidence exists regarding the role of social aspects of tutor presence in juvenile female sensory song learning^6,23^. Additionally, constitutive activation of mTOR during tutor sessions in juvenile males reduced their engagement with the tutor^13^. We asked if our pharmacological manipulations similarly affected the social interactions of the juvenile females with the tutor. Because the juvenile and tutor were in separate cages during tutor sessions, we quantified LookTime—the proportion of time juveniles spent in the region of the cage adjacent to the tutor while facing the tutor—as a proxy for social engagement. We then analyzed LookTime across groups using a Linear Mixed-Effects Model. No significant main differences were observed between the seven groups in the LookTimes (F (5,93) = 0.85, p=0.52). We also did not observe any significant main effect of the day of tutor session (F (2,93) = 1.3, p=0.25) or interaction between groups and days (F (10,93) = 0.82, p=0.605) (**Fig. 7**).

**Figure 7:**
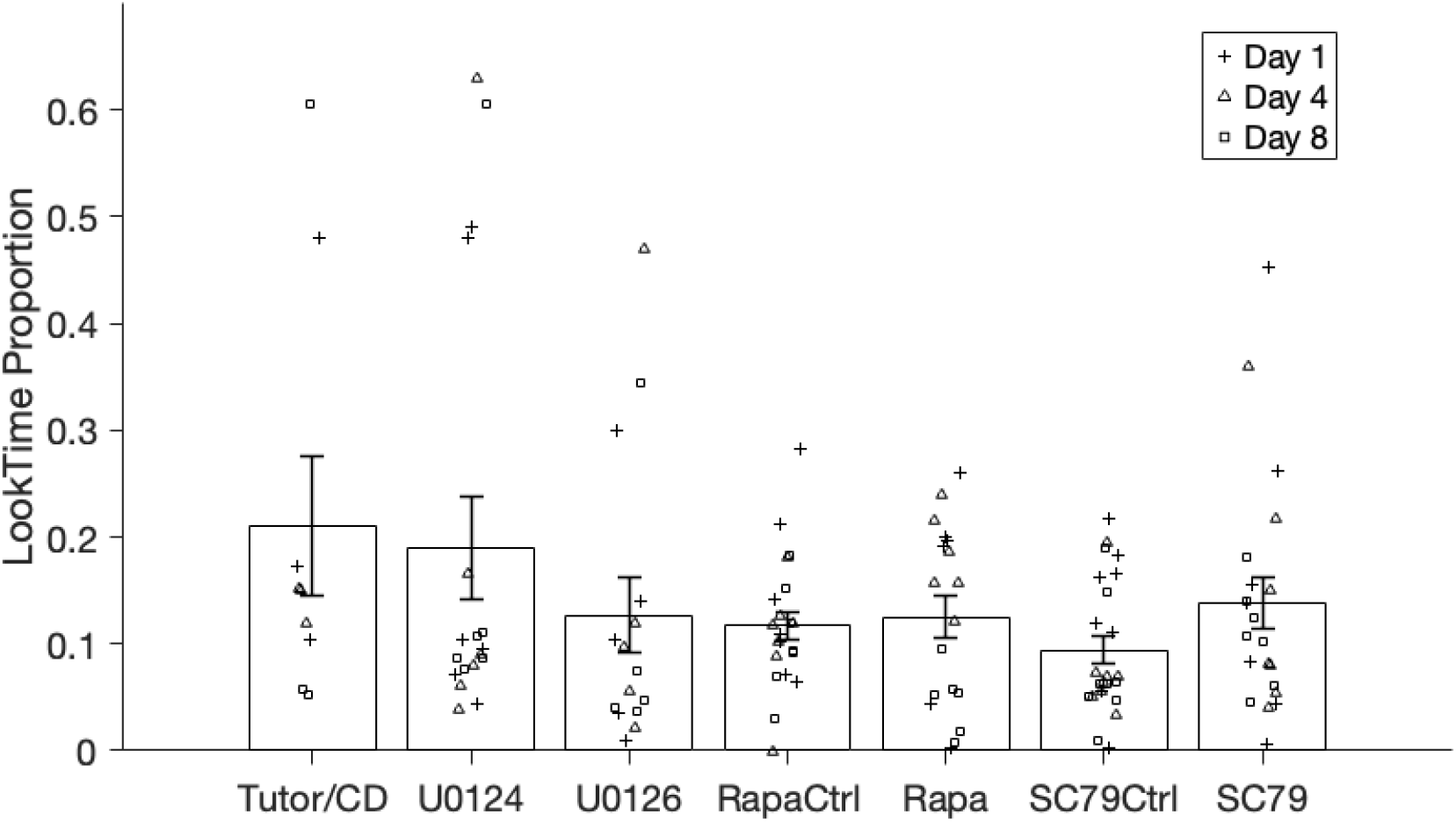
No significant difference between the groups was observed for the times spent in the HZ facing the tutor bird. Bars show means, error bars denote SEMs.

## DISCUSSION

Female zebra finches undergo developmental sensory song learning: exposure to song during development biases adult song preferences toward the songs they heard, whereas females deprived of song experience fail to form consistent song preferences^5,14,24–26^. The adult auditory forebrain in females is genomically and molecularly responsive to song stimuli^27–29^. Song stimuli also induce immediate early gene induction in the female juvenile auditory forebrain^17,27,28^. However, the neuroanatomical locus or molecular mechanisms supporting developmental song learning had not been functionally tested in females. Using pharmacological manipulation of the ERK and mTOR signaling pathways during controlled tutoring, together with an operant perch-trigger assay for adult song preference, we demonstrate that: (1) the female auditory forebrain is causally required for developmental song learning; (2) ERK signaling within the female auditory forebrain is necessary for the formation of song preferences; and (3) the role of mTOR signaling in developmental song learning may differ between males and females, pointing towards at least one sex-specific molecular mechanism underlying developmental song learning.

ERK activation links neuronal activity to long-lasting changes in gene expression, synaptic plasticity, and memory consolidation^30–32^. ERK-dependent induction of ZENK has been implicated in experience-dependent plasticity in the context of song recognition and auditory learning^12,33,34^. ERK signaling in the AL is necessary for song learning in male zebra finches^12^. Given that song-induced ZENK activation^17,35^ as well as the developmental timeline of song learning are largely shared between the sexes^8,9,25^, we asked whether ERK signaling is similarly involved in female developmental song learning. To test this, we infused U0126 — a potent, specific, and transient inhibitor of mitogen-activated protein kinase/ERK kinase^36^ — into the auditory forebrain during developmental song exposure to inhibit ERK signaling. Inhibition of ERK activation during developmental song exposure in females disrupted preference for the tutor song in adulthood, with the preferences of the U0126 group being comparable to that of the Isolate group. This demonstrates that ERK signaling in the auditory forebrain is necessary for developmental sensory song learning in females, as it is in males. Importantly, we provide the first functional evidence that the female auditory forebrain is necessary for developmental song learning.

While ERK signaling appears to be similarly required in both sexes, our data reveal that this parallel does not extend to mTOR-dependent translation. The mTOR cascade regulates activity dependent protein synthesis via its downstream effector S6; phosphorylated S6 (pS6) is needed for functional ribosomes^37,38^. pS6 is thus a marker for *de novo* protein synthesis and therefore active learning via the mTOR-S6K axis^39^. Activity-dependent protein synthesis is an established requirement for long-term memory formation, and developmental sensory song learning represents one such form of long-term memory. The mTOR machinery is present in juvenile females^13^. However, here we found that the Rapamycin group preferred the tutor song significantly over the unfamiliar song much like the Tutor/CD group, indicating that mTOR inhibition did not disrupt developmental song learning in females. This is in contrast with what is observed in males, where inhibition of mTOR disrupts sensory song learning^13^. An explanation for this difference could be that tutor experience might not recruit mTOR signaling in females. Consistent with this idea, song does not induce phosphorylation of S6 in females — at P23, P30, P45, or P60^13,18^ — the developmentally sensitive window for song learning, but does at so at P67^18^, after song preference learning has largely ended^10,14^. In males, however, song-induced phosphorylation of S6 starts at P30 – the opening of the CP, and continues throughout development, until at least P67^13,18^. Protein synthesis supporting long term memory formation for developmental song learning in female juvenile auditory forebrain might thus occur independently of the mTOR-pS6 pathway, using a mechanism distinct from that in males.

We next tested the effect of constitutive activation of the mTOR pathway. Because no specific activator for mTOR exists, we instead used SC79. SC79 is a constitutive activator of AKT and AKT is upstream of mTOR^37^, thus activating AKT indirectly activates mTOR. We found that the SC79 group did not demonstrate a significant tutor song preference and their preference phenotypes were comparable to that of the Isolate group. This mirrors the observation in males where constitutive activation of the mTOR pathway using SC79 disrupts high fidelity song copying^13^. One possibility is that the constitutive activation of mTOR is more disruptive than its inhibition in females. This, however, is difficult to reconcile with the lack of song-induced pS6 activation (discussed above). Given that AKT mediates a boarder range of signaling cascades^40^ beyond just the mTOR pathway, the effect observed with the SC79 group(s) could reflect the involvement of one or more of these alternative SC79 branches.

Further, to separate the contribution of mTOR signaling pathway to stimulus encoding from its contribution to memory consolidation, we included the RapaCtrl and SC79Ctrl groups. These birds received infusions offset from the tutor-song experience. Interestingly, neither the RapaCtrl nor the SC79Ctrl groups demonstrated significant tutor preference. Their preference phenotypes were comparable to that of the Isolate birds. Taken together with the results from the Rapamycin and SC79 groups, these results present a complex picture. SC79 has an effective half-life of approximately 3 hours in the auditory forebrain^13^. Infusion 30 min before tutor exposure, as in the SC79 group, would therefore primarily affect encoding processes. SC79Ctrl birds however, received the drug 2 hours before half of the tutoring sessions and 2 hours after the remaining half. While this would primarily hit the memory consolidation window, it would also result in some overlap with stimulus encoding period. Nevertheless, the disruption of adult tutor preference in both the SC79 and SC79Ctrl groups suggests that constitutive AKT activation is detrimental for both encoding and consolidation in females. Interestingly again, temporally offset infusions of SC79 in males does not disrupt learning^13^, indicating that the disruptive effects of AKT activation in females, unlike in males, might extend to both stimulus encoding and consolidation processes.

To interpret the results of the RapaCtrl group, the longer effective half-life of rapamycin in the auditory forebrain of around 15 hours must be considered^13^. Birds in the RapaCtrl group received infusions 4 hours after each tutoring session, resulting in drug activity during consolidation but offset from tutor experience. However, rapamycin administered 30 min before the 1.5 h tutoring sessions in the Rapa group would also have remained active throughout the post-tutoring consolidation period. Thus, the intact learning observed in the Rapa group and the disruption observed in the RapaCtrl group are hard to interpret. Again, this pattern differs from that observed in males, where song learning was not disrupted in the RapaCtrl birds^13^. Understanding the mechanisms underlying these findings will selective manipulation of additional signaling nodes in the AKT/mTOR pathway.

Importantly, all groups consistently exhibited a significant preference for the song perches over the silent control perches. Our pharmacological manipulations thus did not impair the reward value of song^29^ or produce deficits in hearing or auditory processing. We also considered the possibility that the disruption in learning observed in the U0126, SC79, RapaCtrl, and SC79Ctrl groups could result from factors associated with the experimental procedure including surgery-induced lesions, handling stress, or the infusion process itself. The U0124 group controlled for these factors. U0124 is an inactive analog of U0126, and thus we expected this group to exhibit tutor song preferences comparable to those of Tutor/CD birds. Consistent with this prediction, the U0124 group showed a significant preference for the tutor song. This, together with the intact learning observed in the Rapa group, indicates that the learning deficits were specific to manipulations of the signaling pathways rather than to the experimental procedures themselves.

Finally, given that song learning is socially mediated^21,22^, and SC79 disrupted engagement in males^13^, we tested if manipulation of ERK and mTOR signaling altered the social engagement of the juvenile females to their tutor. We found that birds across all groups engaged with the tutor to a comparable extent. The effects of ERK and mTOR pathway manipulations on adult song preferences in females are thus unlikely to be explained by differences in social attention or motivation during tutoring.

Our study establishes the female auditory forebrain as a causal locus for developmental sensory song learning. We also reveal both conserved and divergent molecular mechanisms underlying developmental learning. While ERK signaling appears to function as a common pathway for song learning in males and females, the complex effects of AKT/mTOR manipulations suggest that not all plasticity mechanisms are recruited similarly across sexes, even though they rely on the same brain area to learn at the same age. Elucidating how signaling pathways are differentially recruited and engaged across sexes and behavioral contexts may provide broader insight into the mechanisms by which early experience shapes the developing brain to have long-lasting behavioral consequences.

## BIBLIOGRAPHY

1. Di Segni M, Andolina D, Ventura R. Long-term effects of early environment on the brain: Lesson from rodent models. Seminars in Cell & Developmental Biology. 2018;77:81–92. doi:10.1016/j.semcdb.2017.09.039

2. Bauman MD. Early Social Experience and Brain Development. J Neurosci. 2006;26(7):1889–1890. doi:10.1523/JNEUROSCI.5133-05.2006

3. Fox SE, Levitt P, Nelson CA. How the Timing and Quality of Early Experiences Influence the Development of Brain Architecture. Child Development. 2010;81(1):28–40. doi:10.1111/j.1467-8624.2009.01380.x

4. Immelmann K. On the effect of early experience upon sexual object fixation in estrildine finches. Published online 1969.

5. Lauay C, Gerlach NM, Adkins-Regan E, DeVoogd TJ. Female zebra finches require early song exposure to prefer high-quality song as adults. Animal Behaviour. 2004;68(6):1249–1255. doi:10.1016/j.anbehav.2003.12.025

6. Riebel K. Early exposure leads to repeatable preferences for male song in female zebra finches. Proc R Soc Lond B. 2000;267(1461):2553–2558. doi:10.1098/rspb.2000.1320

7. Roper A, Zann R. The onset of song learning and song tutor selection in fledgling zebra finches. Ethology. 2006;112(5):458–470. doi:10.1111/j.1439-0310.2005.01169.x

8. Immelmann K. Song Development in Zebra Finch and Other Estrildid Finches. Published online 1967.

9. Eales LA. Song learning in zebra finches: some effects of song model availability on what is learnt and when. Animal Behaviour. 1985;33(4):1293–1300. doi:10.1016/S0003-3472(85)80189-5

10. Riebel K. Developmental influences on auditory perception in female zebra finches - is there a sensitive phase for song preference learning? Animal Biol. 2003;53(2):73–87. doi:10.1163/157075603769700304

11. Riebel K, Smallegange IM, Terpstra NJ, Bolhuis JJ. Sexual equality in zebra finch song preference: evidence for a dissociation between song recognition and production learning. Proc R Soc Lond B. 2002;269(1492):729–733. doi:10.1098/rspb.2001.1930

12. London SE, Clayton DF. Functional identification of sensory mechanisms required for developmental song learning. Nat Neurosci. 2008;11(5):579–586. doi:10.1038/nn.2103

13. Ahmadiantehrani S, London SE. Bidirectional manipulation of mTOR signaling disrupts socially mediated vocal learning in juvenile songbirds. Proc Natl Acad Sci USA. 2017;114(35):9463–9468. doi:10.1073/pnas.1701829114

14. Clayton NS. Song discrimination learning in zebra finches. Animal Behaviour. 1988;36(4):1016–1024. doi:10.1016/S0003-3472(88)80061-7

15. Braaten RF. Song recognition in zebra finches: Are there sensitive periods for song memorization? Learning and Motivation. 2010;41(3):202–212. doi:10.1016/j.lmot.2010.04.005

16. Diez A, Cui A, MacDougall-Shackleton SA. The neural response of female zebra finches (Taeniopygia guttata) to conspecific, heterospecific, and isolate song depends on early-life song exposure. Behavioural Processes. 2019;163:37–44. doi:10.1016/j.beproc.2017.12.022

17. Tomaszycki ML, Sluzas EM, Sundberg KA, Newman SW, DeVoogd TJ. Immediate early gene (ZENK) responses to song in juvenile female and male zebra finches: Effects of rearing environment. J Neurobiol. 2006;66(11):1175–1182. doi:10.1002/neu.20275

18. Butler RM, Delis KD, Maheshwar KV, Montañez KD, Yalcindag S, London SE. Molecular marker of memory formation reveals complex mechanisms of developmental learning. BMC Neurosci. Published online May 28, 2026. doi:10.1186/s12868-026-01013-6

19. Davis HP, Squire LR. Protein synthesis and memory: a review. Psychol Bull. 1984;96(3):518–559.

20. Stankowich T. Quantifying Behavior the JWatcher Way. Daniel T. Blumstein and Janice C. Daniel. Integrative and Comparative Biology. 2008;48(3):437–439. doi:10.1093/icb/icn005

21. Chen Y, Matheson LE, Sakata JT. Mechanisms underlying the social enhancement of vocal learning in songbirds. Proc Natl Acad Sci USA. 2016;113(24):6641–6646. doi:10.1073/pnas.1522306113

22. Derégnaucourt S, Poirier C, Kant AVD, Linden AVD, Gahr M. Comparisons of different methods to train a young zebra finch (Taeniopygia guttata) to learn a song. Journal of Physiology-Paris. 2013;107(3):210–218. doi:10.1016/j.jphysparis.2012.08.003

23. Wall EM, Woolley SC. Social experiences shape song preference learning independently of developmental exposure to song. Proc R Soc B. 2024;291(2024):20240358. doi:10.1098/rspb.2024.0358

24. Miller DB. The acoustic basis of mate recognition by female Zebra finches (Taeniopygia guttata). Animal Behaviour. 1979;27:376–380. doi:10.1016/0003-3472(79)90172-6

25. Riebel K. Chapter 6 Song and Female Mate Choice in Zebra Finches: A Review. In: Advances in the Study of Behavior. Vol 40. Elsevier; 2009:197–238. doi:10.1016/S0065-3454(09)40006-8

26. Williams H, Kilander K, Sotanski ML. Untutored song, reproductive success and song learning. Animal Behaviour.

27. Bailey DJ, Rosebush JC, Wade J. The hippocampus and caudomedial neostriatum show selective responsiveness to conspecific song in the female zebra finch. J Neurobiol. 2002;52(1):43–51. doi:10.1002/neu.10070

28. Bailey DJ, Wade J. Differential expression of the immediate early genes FOS and ZENK following auditory stimulation in the juvenile male and female zebra finch. Molecular Brain Research. 2003;116(1-2):147–154. doi:10.1016/S0169-328X(03)00288-2

29. Lin LC, Vanier DR, London SE. Social Information Embedded in Vocalizations Induces Neurogenomic and Behavioral Responses. Bolhuis JJ, ed. PLoS ONE. 2014;9(11):e112905. doi:10.1371/journal.pone.0112905

30. Thiels E, Klann E. Extracellular Signal-Regulated Kinase, Synaptic Plasticity, and Memory. Reviews in the Neurosciences. 2001;12(4). doi:10.1515/REVNEURO.2001.12.4.327

31. Ojea Ramos S, Feld M, Fustiñana MS. Contributions of extracellular-signal regulated kinase 1/2 activity to the memory trace. Front Mol Neurosci. 2022;15:988790. doi:10.3389/fnmol.2022.988790

32. Impey S, Obrietan K, Storm DR. Making New Connections. Neuron. 1999;23(1):11–14. doi:10.1016/S0896-6273(00)80747-3

33. Cheng HY, Clayton DF. Activation and Habituation of Extracellular Signal-Regulated Kinase Phosphorylation in Zebra Finch Auditory Forebrain during Song Presentation. J Neurosci. 2004;24(34):7503–7513. doi:10.1523/JNEUROSCI.1405-04.2004

34. Jin H, Clayton DF. Localized Changes in Immediate-Early Gene Regulation during Sensory and Motor Learning in Zebra Finches. Neuron. 1997;19(5):1049–1059. doi:10.1016/S0896-6273(00)80396-7

35. Scully EN, Hahn AH, Campbell KA, McMillan N, Congdon JV, Sturdy CB. ZENK expression following conspecific and heterospecific playback in the zebra finch auditory forebrain. Behavioural Brain Research. 2017;331:151–158. doi:10.1016/j.bbr.2017.05.023

36. Favata MF, Horiuchi KY, Manos EJ, et al. Identification of a Novel Inhibitor of Mitogen-activated Protein Kinase Kinase. Journal of Biological Chemistry. 1998;273(29):18623–18632. doi:10.1074/jbc.273.29.18623

37. Hay N, Sonenberg N. Upstream and downstream of mTOR. Genes Dev. 2004;18(16):1926–1945. doi:10.1101/gad.1212704

38. Graber TE, McCamphill PK, Sossin WS. A recollection of mTOR signaling in learning and memory. Learn Mem. 2013;20(10):518–530. doi:10.1101/lm.027664.112

39. Knight ZA, Tan K, Birsoy K, et al. Molecular Profiling of Activated Neurons by Phosphorylated Ribosome Capture. Cell. 2012;151(5):1126–1137. doi:10.1016/j.cell.2012.10.039

40. Manning BD, Toker A. AKT/PKB Signaling: Navigating the Network. Cell. 2017;169(3):381–405. doi:10.1016/j.cell.2017.04.001

